# Explicit strategies in force field adaptation

**DOI:** 10.1101/694430

**Authors:** Raphael Schween, Samuel D. McDougle, Mathias Hegele, Jordan A. Taylor

## Abstract

In recent years, it has become increasingly clear that a number of learning processes are at play in visuomotor adaptation tasks. In addition to the presumed formation of an internal model of the perturbation, learners can also develop explicit knowledge allowing them to select better actions in responding to a given perturbation. Advances in visuomotor rotation experiments have underscored the important role that such “explicit learning” plays in shaping adaptation to kinematic perturbations. Yet, in adaptation to dynamic perturbations, its contribution has been largely overlooked, potentially because compensation of a viscous force field, for instance, is difficult to assess by commonly-used verbalization-based approaches. We therefore sought to assess the contribution of explicit learning in learners adapting to a dynamic perturbation by two novel modifications of a force field experiment. First, via an elimination approach, we asked learners to abandon any cognitive strategy before selected force channel trials to expose consciously accessible parts of overall learning. Learners indeed reduced compensatory force compared to standard Catch channels. Second, via a manual reporting approach, we instructed a group of learners to mimic their right hand’s adaptation by moving with their naïve left hand. While a control group displayed negligible left-hand force compensation, the Mimic group reported forces that approximated right-hand adaptation but appeared to under-report the velocity component of the force field in favor of a more position-based component. We take these results to clearly demonstrate the contribution of explicit learning to force adaptation, underscoring its relevance to motor learning in general.

**New & Noteworthy:** While the role of explicit learning has recently been appreciated in visuomotor adaptation tasks, their contribution to force field adaptation has not been as widely acknowledged. To address this issue, we employed two novel methods to assay explicit learning in force field adaptation tasks and found that learners can voluntarily control aspects of force production and manually report them with their untrained limb. This suggests that an explicit component contributes to force field adaptation and may provide alternative explanations to behavioral phenomena commonly thought to reveal a complex organization of internal models in the brain.

## Introduction

Sensorimotor adaptation is considered important for maintaining skilled motor performance and has been studied extensively, with adaptation to kinematic (Helmholtz 1867; Cunningham 1989) and dynamic sensorimotor perturbations (Lackner and Dizio 1994; Shadmehr and Mussa-Ivaldi 1994) serving as model paradigms. When imposed perturbations are removed after training, an adapted visuomotor map is evidenced by characteristic aftereffects. The transfer of these aftereffects to non-practiced situations has been taken as evidence that the brain learns an internal model, which mimics the input-output relationship of the system being controlled (Shadmehr and Mussa-Ivaldi 1994; Wolpert and Kawato 1998), and uses them for anticipatory compensation of expected perturbations. The observation that these aftereffects sometimes occur despite the learners’ awareness that the perturbation has been removed has been taken as evidence that adaptation of internal models is an implicit process, inaccessible to awareness or intentional control. Furthermore, the fact that aftereffects in amnesic patients, including patient HM, persisted over days has indicated that adaptation does not depend on declarative memory (Shadmehr et al. 1998).

Nevertheless, it has also been shown that learners can leverage declarative or propositional knowledge (Stanley and Krakauer 2013), to consciously modify actions in the face of a perturbation (Cohen 1967; Jakobson and Goodale 1989; Redding and Wallace 2002; Heuer and Hegele 2008; Taylor and Ivry 2011; Taylor et al. 2014), a capacity that we refer to as “explicit learning.” Recent advances in visuomotor rotation paradigms have quantified this component’s contribution to kinematic perturbations: Heuer and Hegele had learners provide perceptual judgements on the movement direction and/or amplitude required to compensate a visuomotor perturbation (Heuer and Hegele 2008, 2009). Benson and colleagues instructed learners about a cursor rotation, using a clock analogy, and tested the quality of their strategy after practice by a structured interview (Benson et al. 2011). Taylor and colleagues asked learners to verbally report their aiming strategy on individual trials based on a set of visual landmarks (Taylor et al. 2014). All of these studies support the idea that explicit learning is a fundamental contributor to the learning curve in such tasks.

Results obtained by these methods suggest that explicit learning may underlie a range of behaviors that were previously explained by extensions of implicit internal models, such as savings (Morehead et al. 2015), structural learning (Bond and Taylor 2017), and context-dependent learning by abstract cues (Hegele and Heuer 2010; Schween et al. 2019), thus highlighting its importance for motor learning (see Krakauer et al. 2019 for a recent review).

For the case of dynamic sensorimotor perturbations, most commonly investigated by perturbing movements through a robot-generated force field (Shadmehr and Mussa-Ivaldi 1994), only a handful of studies have addressed potential explicit learning (Kurtzer et al. 2003; Hwang et al. 2006; Keisler and Shadmehr 2010; McDougle et al. 2015; Thürer et al. 2016). Kurtzer and colleagues instructed a group of participants to “match the effort” of their baseline movements when confronted with a force field and found that it diminished learning compared to a group instructed to “match the kinematics” (Kurtzer et al. 2003). This disengagement by instruction seems incompatible with implicit adaptation in visual perturbation experiments, which proceeds so stereotypically that it can sometimes harm task performance (Mazzoni and Krakauer 2006; Taylor and Ivry 2011; Schween et al. 2014; Lee et al. 2018a). Keisler and Shadmehr, assuming a two-process model with a “fast” and a “slow” component of force field learning (Smith et al. 2006), found that a declarative memory task retroactively interfered with the fast component (Keisler and Shadmehr 2010), suggesting that this fast component, being susceptible to declarative memory load, might be explicit in nature (Morehead et al. 2011). This is further supported by McDougle and colleagues, who asked learners to report their off-target aim in response to a force field using a circular array of visual landmarks. The authors found that the time course of aiming reports overlapped with the putative fast process of adaptation (McDougle et al. 2015).

Despite these clues, the role of explicit learning in force field adaptation still appears to be considered very minor, if not negligible. A factor contributing to this may be that explicit learning has often been linked to a capacity for verbal report, and learners in force field paradigms are frequently unable to report an explicit strategy or even describe the force field’s dynamics when asked to do so. However, the ability to verbalize is limited by the learner’s capacity for communication, and thus may not adequately assess explicit learning in all cases (Stadler 1997; Stanley and Krakauer 2013). It seems reasonable to assume that responses to dynamic force fields are less easily verbalized than simple off-target aiming in response to cursor rotations, and this may explain why learners are unable to adequately report strategies in force field experiments. Previous attempts to ease this problem may be considered limited because these either assessed awareness as an indirect marker of explicit learning (Hwang et al. 2006), or used assessment techniques, such as a circular array of landmarks for reporting aiming angles, that do not reflect the relevant task dimension (Sing et al. 2009; McDougle et al. 2015).

We therefore revisited the question of explicit learning contributions to force field adaptation using two novel experimental paradigms designed to minimize the problems surrounding verbalization. Our first approach (experiments 1 and 2) was rooted in “elimination” techniques frequently utilized in visuomotor rotation paradigms (Heuer and Hegele 2008; Werner et al. 2015): We circumvented the problem of learners verbally describing a complex strategy by instead instructing them to refrain from using strategies on selected force channel trials (Scheidt et al. 2000). Our second approach (experiment 2) aimed to obtain reports in a dimension more suited to the viscous force field than verbal reports (McDougle et al. 2015): Here, we asked participants to mimic the force compensation learned with their right hand using their left hand, allowing us to measure the relevant dimension (e.g., velocity, position) of explicit force compensation strategies.

## Methods

We recruited 87 human volunteers from the participant pool maintained by Princeton University’s Psychology Department to participate in the experiments in exchange for payment or course credit. All participants provided written, informed consent. Experimental protocols were approved by Princeton University’s Institutional Review Board and complied with the relevant guidelines and regulations.

The apparatus was a Kinarm End-Point Lab (RRID: SCR_017060) run with commercial software (Dexterit-E) in a unimanual (experiments 1 and 2) or bimanual configuration (experiment 3). Participants sat in front of the robotic manipulandum, grasping its handle(s) at approximately the level of the lower chest. They rested their head against the edge of a downward-facing horizontal 47-inch, 1920×1080 pixel resolution LCD monitor (LG47LD452C, LG Electronics), and gazed into a horizontal mirror mounted below the monitor. This configuration creates the illusion that the visual display appears in the plane of the movement. By moving the robot handle, participants could move a screen cursor (blue disc, 10 mm diameter) that matched handle position.

The robot could generate field, null, and channel trials. On field trials, the robot generated a velocity-dependent force field (Shadmehr and Mussa-Ivaldi 1994): 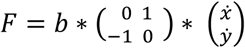, where 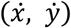 are directional velocities and a positive field constant *b* creates a clockwise force field. On null trials *b* was zero, and the robot did not actively influence the movement. On channel trials (Scheidt et al. 2000), the robot constrained the movement to a straight line through start and target by generating a spring of 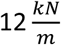 with a damping coefficient of 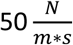 (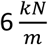 and 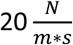 for experiment 3). The channels served as test trials, allowing us to measure the predictive forces participants learned to exert in response to the field in the absence of stiffness influences.

### Experiment 1

Each trial began with the robot passively moving the participant to a start location (white disc, 15 mm diameter) on the participant’s midline, approx. 45cm in front of their chest. After the participant held this position for 200 ms, a target (white disc, 15 mm diameter) appeared on the midline, 120 mm from the start position, and the participant’s task was to slice through the target in a rapid shooting movement. We chose these open-loop movements rather than more standard closed-loop movements as our channel instructions (see below) would theoretically have prevented task success on field trials, and we did not want our participants to reason about the difference between field and channel trials. Target hits were reinforced by a pleasant “ding” sound. Movement speed was incentivized by a “too slow” message appearing if movement time exceeded 350 ms. If participants moved before the appearance of the target or if movement time exceeded 1000 ms, the trial was aborted and repeated.

Twenty-five participants (mean age: 21, range: 18-26 years, 16 female) practiced a force field with 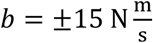 and we tested their predictive force compensation on channel trials. Out of these, three participants were not analyzed due to technical difficulties, and two more participants were excluded because standardized post-experimental questioning indicated that they failed to follow instructions. It was critical in this experiment for participants to follow instructions because we sought to test if they could intentionally control part of the adaptive response to the force field. This required them to implement the correct intention on the respective channel trial. Responses that led to exclusion included indicating that they “still pushed” on “No Push” trials (see below) or that they “pushed hard straight ahead” (i.e. towards the target) on Push trials.

We introduced two new types of channel trials in addition to the standard Catch channel: On these trials, participants were provided with an onscreen message prior to the reach (see figure 1A for experiment 2 methods, which are similar), reminding them of an instruction given approximately 10 trials before the force field was introduced: One message read, “No Push”. For this message, participants were instructed to “act as in trials where the robot doesn’t push you off path and just move towards the target”. We thus aimed to probe their ability to voluntarily disengage a hypothetical explicit strategy. The other message read “Push”, reminding participants of the instruction to “expect the robot to push you off and act as on those trials”. We included this “Push” trial type as a control to ensure that any force modulation would be attributable to the associated instruction rather than other factors, such as delays introduced by the messages allowing a labile component of implicit memory to decay (Miyamoto et al. 2014; Zhou et al. 2017). Standard Catch channel trials were not preceded by a message. The full, scripted instructions (of experiment 2) are provided in supplementary text 1.

**Figure 1:**
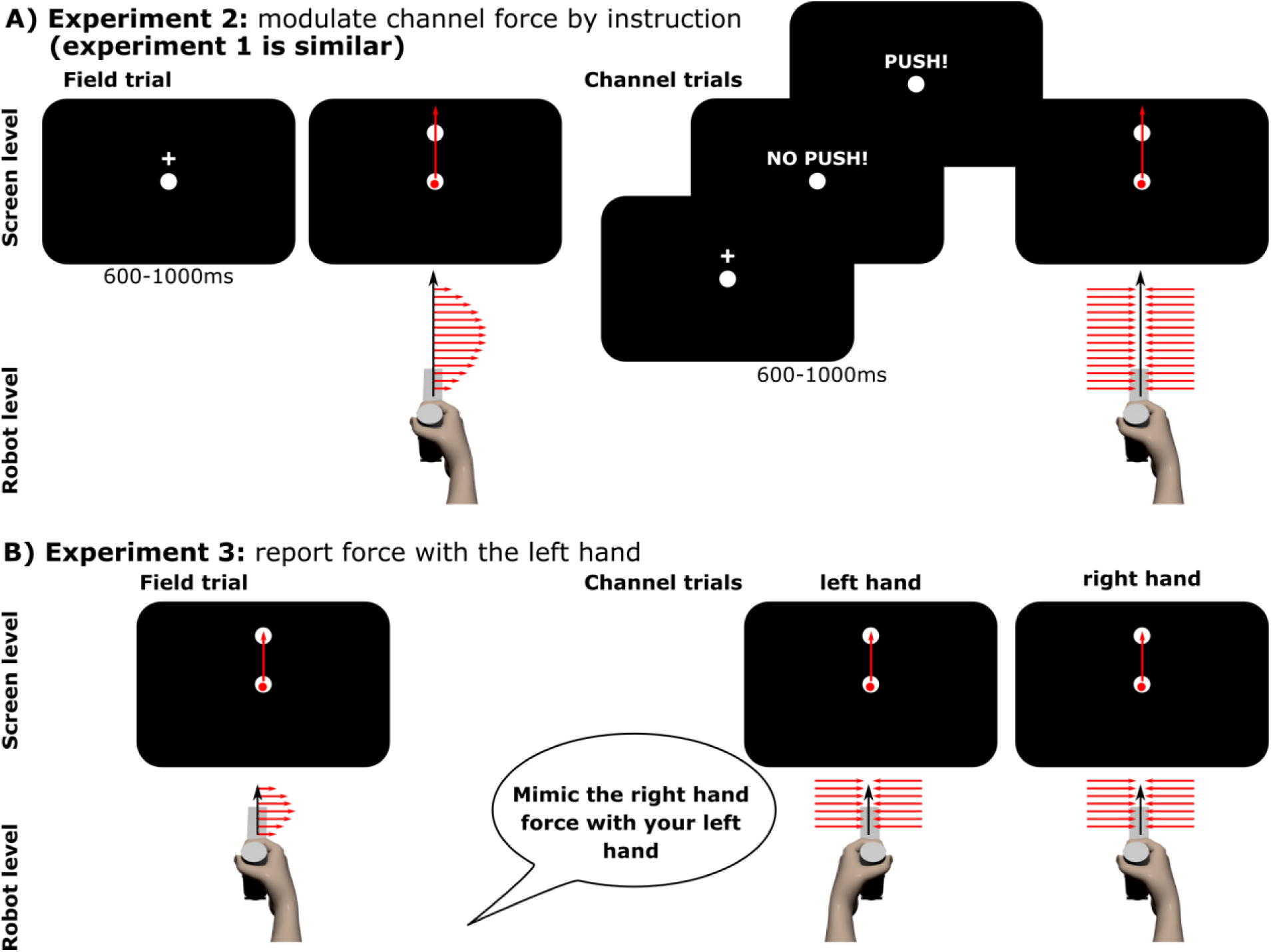
Schematic methods. A) Experiment 2 (experiment 1 is similar): On field trials, participants saw a fixation cross after the robot had guided them back to the start location. When the cross disappeared and the target appeared, their task was to “shoot through” the target in a single, fast movement, experiencing the force field (or null field in baseline). The red arrows schematically illustrate (potential) robot-generated forces. Note that the movement path on field trials could deviate substantially from the straight line illustrated. On channel trials (Scheidt et al. 2000), the robot constrained movements to the line connecting start and target. Catch channels were preceded by the fixation cross, making them indistinguishable, a priori, from field trials. On instructed channels, instead of the fixation cross, learners would see a keyword prompting them to either expect the robot not to push them off their path or to do so, and act accordingly. B) Experiment 3: Participants practiced closed-loop target reaches in the field with their right hand and were tested on right hand (Catch) and left hand channels. On left hand channels, the Mimic group was instructed to mimic their right hand forces. The control group received no such instruction.

The protocol was an A-B-Clamp paradigm (Smith et al. 2006; McDougle et al. 2015), consisting of 40 trials familiarization, 60 trials baseline, 450 trials field A practice, 45 trials field B practice and 50 trials clamp. In the familiarization phase participants moved from start to target in a null field to get acquainted to the task and the time criterion. Baseline was also in a null field, but contained six Catch channels to establish a baseline, with one channel per block of ten trials and an additional constraint that channels could not be the first or last trial of a block, ensuring there were at least two non-channel trials between any two channels. In field A practice, a force field was introduced, with the sign counterbalanced across participants. Each block of ten trials contained one channel, as before (i.e. 10% channels), but now, metablocks of three blocks contained each of the three channel types (Catch, No Push, Push) once, in a random order. Field A practice thus contained 15 channels per type. Field B practice exposed participants to a force field of opposite polarity than field A. This phase contained one channel per block of five trials (i.e. 20% channels), with all other constraints as in field A. The final clamp phase consisted of only channel trials. This block no longer had screen messages, but participants were verbally instructed to consider all trials No Push trials from the beginning of the clamp phase until the end of the experiment to reveal behavior in the absence of potential explicit strategies (McDougle et al. 2015). We used the same randomization sequence for all of our participants in experiment 1 and 2.

### Experiment 2

Experiment 2 was designed to test whether behavior on No Push channels would match behavior on a more commonly used, longer error clamp period with the instruction that the perturbation was removed. We therefore omitted field B practice and instead had participants proceed directly into the final clamp phase, which was 95 trials long. Furthermore, the experiment served as a replication of the main effect between channel types in experiment 1.

We tested 22 participants (mean age: 21, range: 18-28 years, 16 female) on this experiment and excluded five because post-experimental questioning indicated that they did not follow the instruction on the Push or No Push channels. (As in experiment 1, the potential success of the experiment was critically dependent on participants following the instructions.)

The procedures were the same as experiment 1 with the following exceptions: we suspected that, in experiment 1, the instructive message tended to startle participants into moving prematurely (resulting in the trial being repeated), because it occurred around the time where they expected the target to appear. To avoid this problem, we now showed participants either the white text message or a white fixation cross (on null, field and Catch trials) after they were in the start for 200 ms and had the text/cross disappear and the target appear after another interval uniformly randomized between 600 and 1000 ms (figure 1A). We considered using another message instead of the fixation cross to match the channel messages in luminance and area, but decided against it as we feared participants may become too accustomed to viewing a message and could potentially ignore it.

### Experiment 3

Experiment 3 aimed to test if participants could mimic their right hand force compensation with their left hand and thereby express their explicit knowledge about the force field in a dimension more suitable to this complex perturbation than directional landmarks (Taylor et al. 2014; McDougle et al. 2015). Participants adapted to a force field by closed-loop movements to a target 100 mm in front of the central start location. Field exposure was interspersed by right hand Catch channels and additional left-hand “Mimic” channels, where they moved with their left instead of their right hand. The Mimic group (N=20, mean age: 21, range: 19-24 years, 17 female, 1 excluded for technical issues) was instructed to “mimic the forces” they were applying with their right hand with the left hand, whereas the control group (N=20, mean age: 22, range: 18-34 years, 11 female, 3 excluded for technical issues) received no such instruction concerning left hand trials. To ensure that all participants could sense the force field, and to more closely match a previous experiment (McDougle et al. 2015), we increased the field constant to 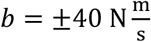. Like experiment 1, the protocol was a field A - field B - Clamp paradigm, now consisting of 100 trials baseline (80 null, 10 right and 10 left hand channel), 200 trials field A practice (158 field, 21 right, 21 left hand channels), 20 trials practice of an opposing field B (18 field, 1 right, 1 left channel) and 100 clamp trials (90 right hand, 10 left hand channels). Field direction was counterbalanced across participants.

### Data analysis

The data were processed in Matlab (RRID: SCR_001622) and analyzed in R (RRID: SCR_001905) and JASP (RRID: SCR_015823) for visualizations and statistics.

For experiment 1 and 2, we defined movement start as the instance when movement speed first exceeded 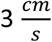 and movement end when the radial distance to start of the hand first exceeded that of the target’s center. In experiment 3, we used a different criterion of 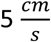 for at least 100 ms, as we noticed the lower criterion included some premature movements whose occurrence we attribute to the different movement types (open vs. closed loop). Movement end in experiment 3 was defined as the hand being within the target circle with a speed of less than 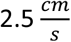. We then truncated force and position data of each movement to the section between these events. We calculated axial velocities using a Savitsky-Golay filter and additional time frames at the start and end of the reach to avoid ramping effects. Data of participants experiencing the CW field in the A-phase were flipped about the y-axis before further processing.

To quantify force compensation on channel trials, we calculated participants’ forces in the channels relative to the ideal force profile that would have perfectly compensated field A, given participants’ velocity along the channel, using zero-offset regression and expressed the slope as percent force compensation. For open-loop movements (experiment 1 and 2), we did this on the complete data from start to end, whereas for closed-loop movements (experiment 3), we used a window of 400 ms prior and posterior to peak speed, or start/end, respectively, if this window exceeded the actual event. These data were aligned to peak speed and padded with NaNs to produce mean trajectories for plotting.

Experiment 3 also measured the degree to which force expression was composed of position and velocity components for the left and right hands. To compute this, we fit perpendicular channel force profiles using a zero-offset linear model of position along the channel, velocity along the channel, or a linear combination of the two, respectively (Sing et al. 2009). We used adjusted R^2^ (as returned by R’s summary.lm function) to quantify variance accounted for, and we averaged across participants by transforming its square root to Fisher-Z-scores (Bortz and Schuster 2010), calculating their mean, and retransforming to R^2^. We did this for 1000 bootstrap samples of individuals in this group, and estimated 95% confidence intervals.

In all experiments, we computed RT on field trials as the time from target display to movement start, and removed RTs exceeding three standard deviations of participants’ individual means (experiment 1: 97 trials, 1 %; experiment 2: 114 trials, 1 %; experiment 3: 154 trials, 1.7 %). We then calculated individual medians over the following blocks: late baseline, early field A practice, late field A practice, field B practice (where applicable) and early clamp, using 36 (experiment 1 and 2) or 18 (experiment 3) field trials per block, and averaged across participants. For experiment 1 and 2, late baseline excluded the last 10 trials of baseline as RTs there were distorted by instructions sometimes being given during the RT interval.

### Statistical analysis

We used mixed ANOVAs with group as between-subject factor, and channel type, hand and/or block as within-subject factors. We checked for unequal variances between groups by Levene’s tests. Where Mauchly’s test indicated significant non-sphericity, we report Greenhouse-Geisser-corrected *p*-values, as discernible from floating point instead of integer degrees of freedom. For effect sizes, we report generalized eta-squared. To follow up on significant effects, we used one-way ANOVAs and t-tests, reporting Bonferroni-Holm corrected *p*-values for the latter. For comparing dependent correlations, we used a test described in (Bortz and Schuster 2010, p. 167-8).

## Results

### Experiment 1

Experiment 1 tested if predictive force compensation in channels is fully implicit, or if it contains an explicit component which learners can voluntarily modulate. Participants practiced open-loop shooting movements and were exposed to two velocity-dependent force fields, where they experienced an initial long block of field A, followed by a short block of an opposing field B, and a clamp block (Smith et al. 2006; McDougle et al. 2015). Channel trials tested participants’ predictive force compensation (Smith et al. 2006) on a fraction of trials. Whereas all baseline channels and one third of channels during practice were standard, unannounced Catch channels, the other two thirds were preceded by an on-screen message instructing participants about the upcoming trial (figure 1A). One message type said “No Push!”, for which our instructions asked participants to “act as in trials where the robot doesn’t push you off path and just move towards the target”. We reasoned that if predictive compensation is fully implicit, participants would express the same amount of force on these trials as they did on Catch trials. On the other hand, if participants could spontaneously downregulate their force in response to the instruction, this would indicate that an explicit component was contributing to Catch channel behavior. The other message type said “Push!”, asking participants to expect the robot to push them off and act accordingly. The B-field was followed by a clamp block of all channel trials (Smith et al. 2006; McDougle et al. 2015), for which we instructed participants that they should treat all of them as “No Push!” trials, even in the absence of messages.

Figure 2A shows the force compensation index across trials for each type of instructed channel. During field A practice, participants expressed less force on No Push compared to Catch channels, revealing an ability to voluntarily eliminate a component of predictive force compensation in response to the instruction. Conversely, channels with the instruction to Push voluntarily show compensation that is not smaller, but even larger than on Catch channels, suggesting that behavior on No Push channels was under volitional control rather than just a general effect of the instruction. Figure 2B visualizes across participant average force profiles on selected channel trials.

**Figure 2:**
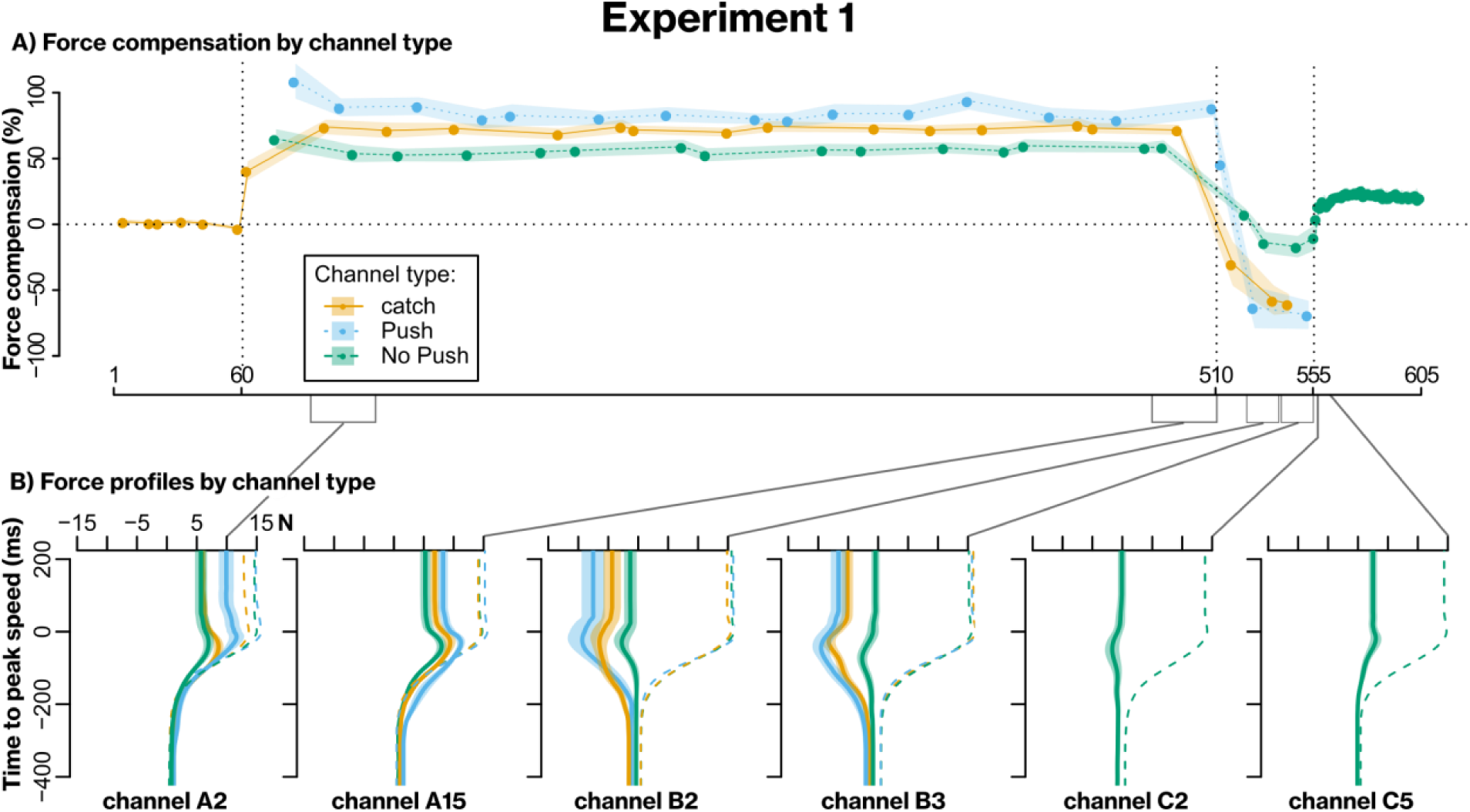
Results of experiment 1. A) Group mean force compensation (±standard error of mean; SEM) across trials. B) Group mean force profiles (±SEM) obtained by averaging individual force profiles aligned to peak speed. Dashed lines are hypothetical, ideal field A compensatory forces calculated from velocity and used as reference. X-Labels reference position in the protocol, with A2 being the 2^nd^ trial in field A, C5 the 5^th^ trial in the clamp phase, etc. Multiple lines per panel are from channels of the different types, respectively. Grey lines and brackets between panels indicate regions represented in this way.

To statistically analyze these data, we averaged the first, middle and last three channels of each type from field A-practice into blocks and performed an ANOVA with the within participant factors channel type and block. This revealed a significant main effect of channel type (*F*_2,36_=25.4, *p*<.001, *η*^*2*^_*gen*_=.27), no significant main effect of block (*F*_1.3,22.7_=.17, *p*=.74, *η*^*2*^_*gen*_=.002), and a significant interaction (*F*_2.0,36.8_=4.7, *p*=.002, *η*^*2*^_*gen*_=.05). Simple main effects indicated that the effect of channel type was present at all three levels of block (early: *F*_*2,36*_=25.3, *p*<.001, *η*^*2*^_*gen*_=.34; middle: *F*_*1.4,25.8*_=15.2, *p*<.001, *η*^*2*^_*gen*_=.31; late: *F*_*1.3,23.5*_=8.9, *p*<.001, *η*^*2*^_*gen*_=.22) and post-hoc *t-*tests at these levels indicated significant differences between all pairs of channel types (all *p*<.02 after Bonferroni-Holm correction), except for the No Push versus Catch channels in the early block (*p*=.21) and for Push versus Catch channels in the late block (*p*=.13). Whereas we might have expected an effect of block, reflecting learning, the time courses of Catch channels (figure 2A) indicate that learning may have been too quick to be captured by the relatively infrequent channel trials. Such quick learning may be explained by the use of a single target, and by increased requirement for predictive compensation imposed by open-loop movements. Regardless, the main effect of channel type suggests that the degree to which participants express learning is under volitional control, which would be consistent with the use of a strategy.

The observation that compensation on the Push channels did not only match, but exceeded Catch channel behavior was unexpected. We assume that the instruction prompted participants to more voluntarily compensate the force than when just expecting a standard field trial, and this could explain the excess force applied. If such a direct effect of the manipulation occurs on Push trials, could it also explain performance on No Push trials? That is, could decreased expression be explained by participants voluntarily pushing in the other direction - which, notably, would still imply them consciously applying force - rather than just stopping to apply an explicit strategy? The time course of behavior when switching to field B does not support this possibility. Specifically, if participants were applying counterforces in direct response to the instruction, we would expect these to remain of approximately the same magnitude in the B-phase, albeit potentially with a sign switch. Whereas this may be seen in the Push channels, behavior on No Push channels appears to follow a time course that is much more independent from that on Catch channels (figure 2A).

Overall, the time courses of the three channel types during field B are consistent with previous findings in this paradigm (McDougle et al. 2015): when participants expect the robot to apply force (Push and Catch channels), predictive compensation quickly changes sign upon exposure to field B, while compensation when they do not expect the robot to apply force (No Push channels) decreases more gradually with practice, so that not only the sign of each individual channel, but also the relation between the three channel types is inverted by the end of the B-phase. This was confirmed by a repeated measures ANOVA on the individual differences between the last channel of field A and field B, respectively, which indicated a significant difference (*F*_1.5,28.2_=12.6, *p*<.001, *η*^*2*^_*gen*_=.40) with post hoc-tests showing a significant difference between No Push and both Catch (*t*=-5.2, *p*<.001) and “Push” trials (*t*=-4.1, *p*=.001), but not between Catch and Push trials (*t*=1.4, *p*=.18). This means that the change from field A to field B was significantly smaller on No Push than Push or Catch trials. Considering the short timeframe in focus, this is in line with the implicit No Push trials capturing a slow component of learning, while an additional fast explicit process is represented in Push and Catch trials (Keisler and Shadmehr 2010; McDougle et al. 2015).

In the final clamp phase consisting of only No Push channels, behavior rebounded. This observation would be difficult to explain by the No Push instruction directly eliciting volitional forces, and is more consistent with behavior seen in implicit learning of force fields and cursor rotations in this paradigm, where it has been explained by a fast, explicit component that is adapted to field B being quickly diminished by instruction or decay, revealing a slow component that is still adapted to field A (Smith et al. 2006; McDougle et al. 2015).

### Experiment 2

Experiment 1 supported our hypothesis that learners can voluntarily disengage a component of channel force compensation when instructed to do so right before the trial. Here, we wanted to verify that this is not just an idiosyncrasy of the single-trial instructions. In experiment 2, we therefore tested if behavior on a more standard, post-experimental clamp phase with the No Push instruction administered only at phase onset, would indeed reflect similar behavior as the interspersed trials and further display the hallmark behavior of gradually decaying memory. We therefore replicated experiment 1, but removed field B practice.

Figure 3 shows that the instruction modulated force compensation on channel trials as before. Accordingly, like for experiment 1, an ANOVA on field A practice with factors block and channel type indicated a significant main effect of channel type (*F*_*2,34*_=, *p*<.001, *η*^*2*^_*gen*_=.16) but not of block (*F*_*1.4,23.4*_=1.3, *p*=.28) and no interaction (*F*_*2.1,35.4*_=.94, *p*=.46). Averaging over blocks and comparing channel types by post-hoc t-tests revealed differences between all pairs of channel types (all *p*<.02 after Bonferroni-Holm correction). Thus, experiment 2 replicates the main effect of channel type observed in experiment 1.

**Figure 3:**
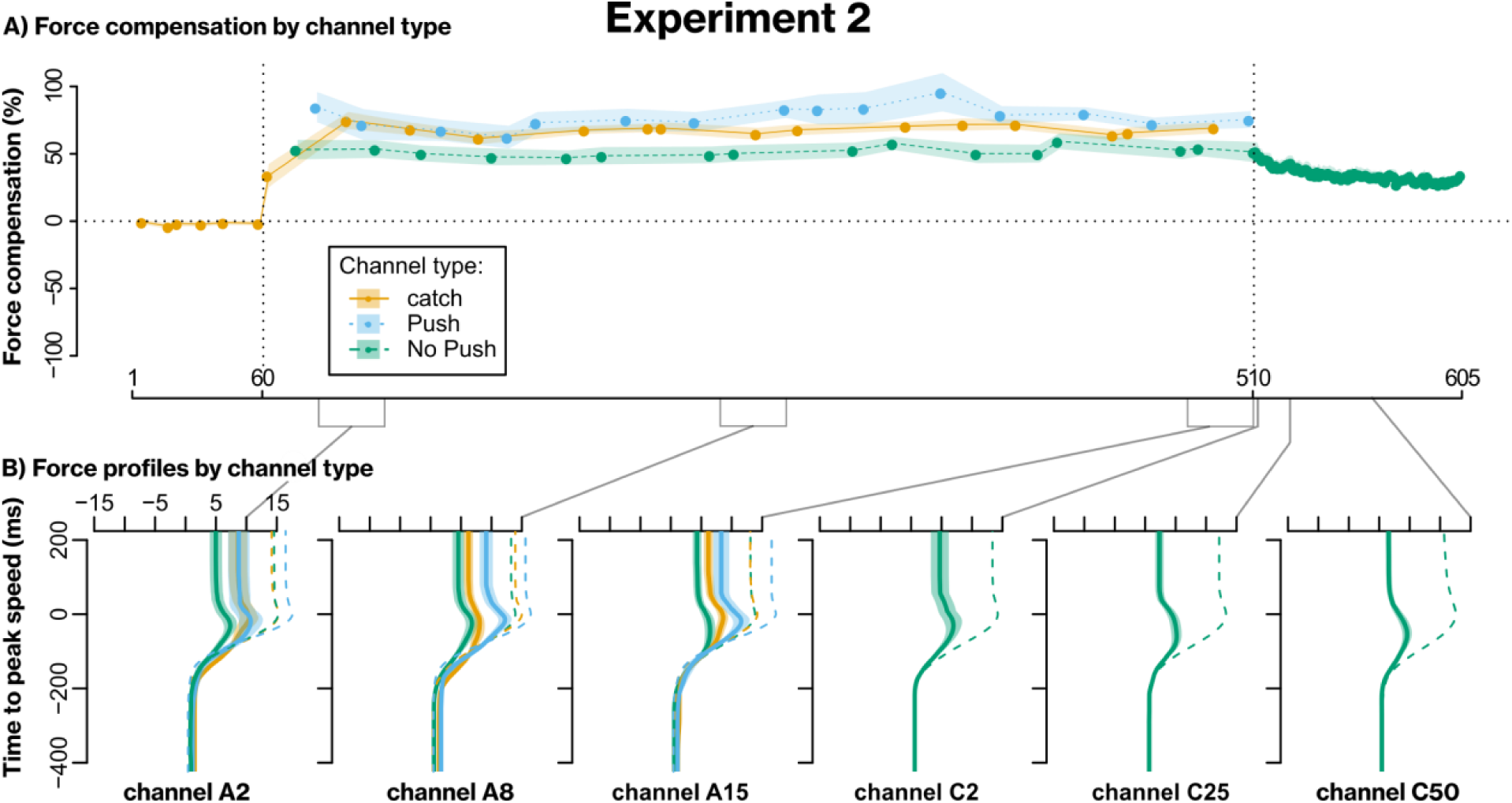
Results of experiment 2. A) Group average force compensation measures across trials. B) Group mean force profiles (±SEM) on selected trials. X-Labels reference position in the experiment, with A2 being the 2^nd^ trial in field A, C25 the 25^th^ trial in the clamp phase, etc.

When going directly from field A into the clamp phase, with the instruction not to push voluntarily, behavior on clamp appeared to pick up on the most recent interspersed No Push channel and to decay smoothly from there. When correlating individual participants’ performance on the first channel of the final phase and on the last interspersed channel of each instruction type, this correlation was significantly stronger for the No Push channel than for the Catch (z = 3.5, *p* < .001) or Push channel (z = 2.9, *p* = .003), and the regression line was close to unity (figure 4). We therefore take this result to indicate that the behavior in the final clamp phase reflects the same implicit processes exposed by the No Push channels interspersed during learning.

**Figure 4:**
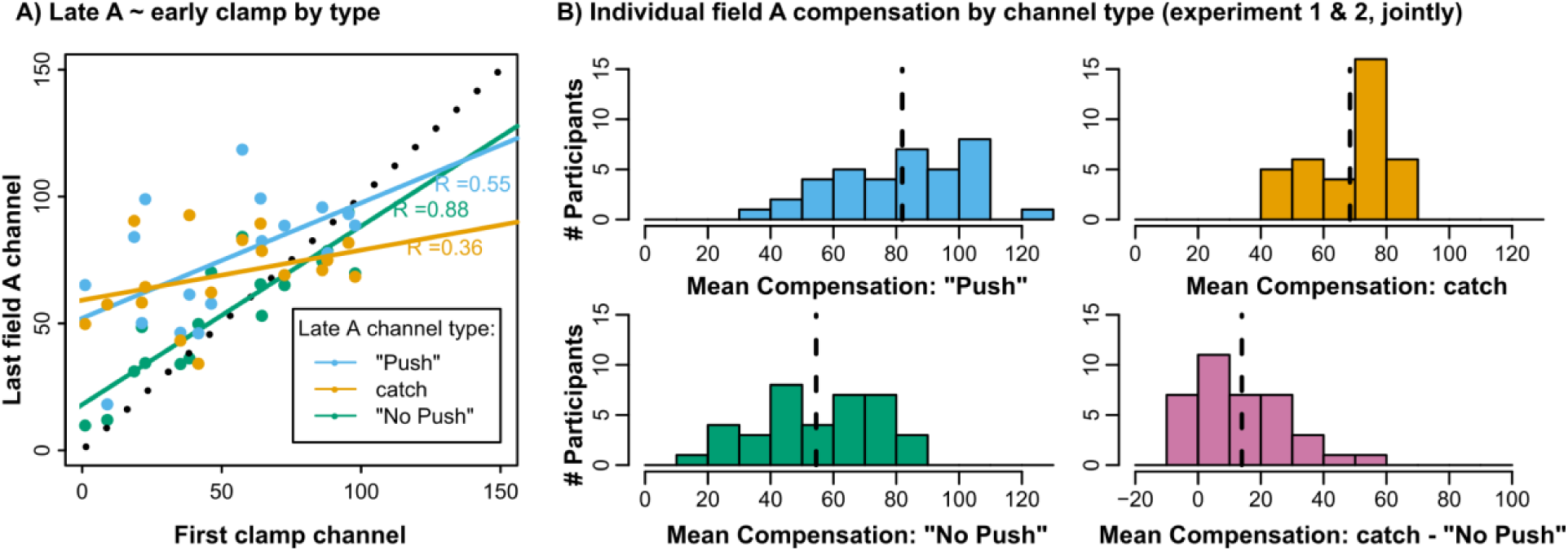
A) Scatterplot with least squares regression lines: last channel of field A practice for the different types against the first clamp channel. Black, dotted line is identity. B) Histograms of individual means of force compensation index across field A practice (joint data from experiment 1 and 2). Dashed, vertical lines mark group means. Panel 4 shows individual differences between No Push and Catch channels.

How is the difference between the channel types reflected in individual participants? Figure 4B shows the histograms of individual average force compensation during field A practice on Catch and No Push channels and the individual differences between the channel types for participants of experiment 1 and 2. It is clear that some participants responded strongly to the instruction whereas others show no force modulation. One explanation for this could be that participants differed in how they understood and followed our instructions. We excluded cases where standardized post-experimental questioning made it clear that participants misunderstood the instructions and either consciously compensated on No Push or did not compensate on “push” channels (see *Methods*). However, some responses were ambiguous so we kept those participants in the analysis to avoid potential selection bias.

Another possibility is that some participants indeed learned more explicitly/less implicitly than others. Large variation between individuals is common in cursor adaptation experiments (Schween and Hegele 2017), and studies have determined that perturbation magnitude relative to baseline movement variability (Oh and Schweighofer 2018; Gaffin-Cahn et al. 2019), as well as participants’ age and specific cognitive abilities (Heuer and Hegele 2008; Anguera et al. 2010, 2011; Hegele and Heuer 2013; Vandevoorde and Orban de Xivry 2018; Wong et al. 2019) are important factors determining if participants become aware of a perturbation, and can develop and apply explicit strategies to counteract it.

In summary, we take results of our manipulation in experiments 1 and 2 to show that predictive compensation of a force field expressed on standard Catch channels was not purely implicit. If learning is truly implicit, we would expect participants to have no volitional control over its expression. Conversely, participants in our experiment were able to modulate predictive expression based on verbal instruction in line with an explicit component contributing to learning.

### Experiment 3

Experiments 1 and 2 show that force compensation performance can be modified based on verbal instruction, thus displaying a key characteristic of explicit learning. In experiment 3, we sought to complement this finding by testing whether participants can express their knowledge about the force field in a task-relevant dimension. In a previous experiment, we presented participants with a circular array of landmarks and asked where they would have to aim their movement to compensate for the perturbation (McDougle et al. 2015). Whereas this reporting method is well suited to cursor rotation experiments, where an angular change in aiming is sufficient to compensate a rotation and the report can therefore be considered to lie on the same scale as actual movement performance, it is potentially not ideal for velocity-dependent force fields. To address this issue, we asked participants to express the movement strategies they were consciously using to compensate for the force field, by mimicking them with their left hand. Based on the finding that true implicit inter-manual transfer of force compensation is limited (Wang and Sainburg 2003; Malfait and Ostry 2004; Joiner et al. 2013), we reasoned that forces expressed with the left hand could thus serve as a proxy of explicit knowledge, which participants had acquired about the force field with their right hand, on a more suitable scale than reporting aiming directions.

We tested two groups of participants on an experiment where the test channels interspersed between closed-loop reaches in a force field were alternatively conducted with the left or right hand by each participant. We asked participants in the Mimic group to “mimic the forces” they experienced on right hand trials with their left hand, whereas the control group was just told to move to the target.

Figure 5A & B show the force compensation index and force profiles of both groups by hand. Both groups compensated the force field about equally with their right hand. As for the left hand, the control group expressed very little force adaptation, confirming our expectation of limited intermanual transfer based on prior results (Wang and Sainburg 2003; Malfait and Ostry 2004; Joiner et al. 2013). For the Mimic group, we noted that different participants appeared to generate forces in different directions (see figure 5B). We interpret this as participants mimicking the force in different reference frames, i.e. in an extrinsic and an intrinsic reference frame. Whereas it has been shown that force compensation can be learned in different and mixed coordinate systems (Berniker et al. 2014; Franklin et al. 2016), it is not clear why implicit acquisition would yield the seemingly categorical differences we observed. If, on the other hand, we assume left hand performance to express participants’ explicit knowledge, the different reference frames are easily explained by differing interpretations of the task instructions: as the instruction did not specify a reference frame, participants chose the one they considered appropriate. We therefore aligned left hand forces in the Mimic group to an extrinsic reference frame by sign-inverting left hand forces of participants whose median reports in field A matched an intrinsic reference frame (8 participants in Mimic group, 4 in control group). With this, left hand force compensation in the Mimic group on average trailed that of the right hand, albeit still with higher variability (figure 5A).

**Figure 5:**
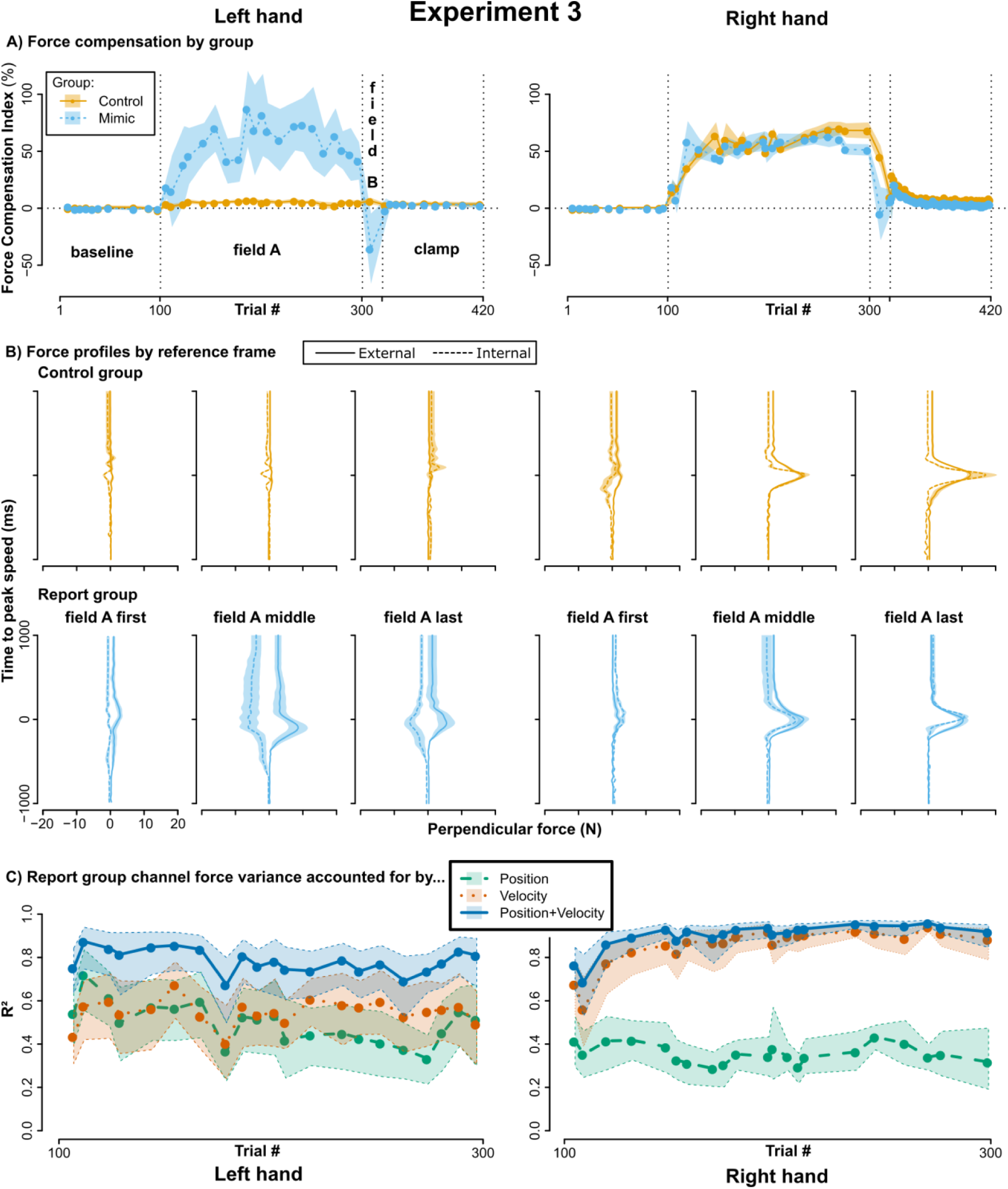
Experiment 3. A) Across participant mean force compensation indices (±SEM) of left and right hand force channel trials for the control group with no instruction and the Mimic group instructed to mimic the forces with their left hand. To average left hand performance, the signs of left hand forces were flipped for 8 out of 19 participants in the Mimic group and 4 out of 17 in the control group, as they appeared to report in an inverted reference frame. B) Mean force profiles (±SEM) at the beginning, middle and end of field A practice, with separate means for participants displaying a more external or more internal reference frame. C) Variance in perpendicular channel force accounted for by along-channel position, velocity, or a linear combination of the two. Shaded areas are bootstrapped 95% confidence intervals.

The group difference was supported by a mixed ANOVA with factors group (control vs. Mimic), block (early, middle, late field A practice) and hand (right, left). This yielded a significant main effect of group (*F*_*1,34*_ = 6.6, *p* = .015, *η*^2^_*gen*_ = .16) and block (*F*_*1.4,48.0*_ = 6.3, *p* = .009, *η*^2^_*gen*_ = .15), a tendency for a hand-effect (*F*_*1,34*_ = 4.1, *p* = .051, *η* ^2^_*gen*_ = .09) and a significant interaction between group and hand (*F*_*1,34*_ = 5.6, *p* = .023, *η*^2^_*gen*_ = .13), but no other significant two- or three-fold interactions (all *p* > .29). Following up on the group-by-hand interaction, simple main effects indicated a significant difference between left and right hand in the control (*F* = 103.8, *p* < .001), but not in the Mimic group (*F* = .04, *p* = .85).

The brevity of the field B phase only allowed us to test one channel per hand in its center. These data were highly variable; for example, the Mimic group had right-hand mean: −5 %, SEM: 101 %, median: 21 %, and left-hand mean: −36 %, SEM: 134 %, median: 5 %). Closer inspection revealed that this was not due to few outliers but indeed reflected the group being spread out between maintaining compensation, switching the sign of their compensation, or even increasing. Accordingly, neither *t*-test (*t*_*18*_=.6, *p*=.55), nor sign-test (*S*=7, *p*=.18) indicated a significant difference between left and right hand for differences between the last field A channel and the field B channel. We speculate that this is reflective of some learners adopting a new strategy that is appropriate to the sign-switched force field, while others did not, potentially because they chose to mimic in a single reference frame. We therefore refrain from overinterpreting the field B or clamp phase in this experiment and note that the method would need to be improved to allow inferences on field switches or rebound.

### Representation of force compensation

Field A practice data afforded us the opportunity to determine if force compensation reflected separable components of learning. If the brain learns an internal model of a velocity-dependent force field, we may expect that the model’s output, i.e. compensatory force, should reflect such a velocity-dependence (Conditt and Mussa-Ivaldi 1999). Indeed, it has been found that the compensatory response to a force field is determined by hand velocity and to a lesser extent by hand position (Sing et al. 2009). The position-dependent component has subsequently been attributed to a different, model-free learning mechanism (Haith et al. 2011). Considering that recent studies have found proposed model-free reinforcement learning to depend on explicit strategies (Shmuelof et al. 2012; Codol et al. 2018; Holland et al. 2018), this can be taken to suggest that the strategic component of learning should depend on position and, in this respect, resemble off-target aiming typical of cursor rotation experiments (Benson et al. 2011; Taylor et al. 2014; Bond and Taylor 2017). Alternatively, strategies could also depend on velocity. Different from previous approaches (McDougle et al. 2015), experiment 3 offered learners a way to express a strategy in exactly the dimension of the field, allowing us to infer the nature of strategy representation. Figure 4D displays the proportion of variance accounted for on all of the field A channels in experiment 3 when fitting channel-perpendicular force by position, velocity, or a linear combination of the two. For the right hand, the proportion of variance accounted for by velocity is larger than that accounted for by position and increases with practice, in line with learning an internal model of the velocity-dependent force field (Sing et al. 2009; Haith et al. 2011). For the left hand, position and velocity accounted for more equal portions of variance throughout practice. Thus, the putative explicit strategy did not appear to capture the velocity-dependent nature of the force field as adequately as right-hand learning, even when the two were assessed by the same method. This supports our interpretation that the explicit component expressed by the left hand is qualitatively different from adaptation of an internal model, and suggests that the complexity of explicit learning may be limited to simple, position-dependent strategies, as are typically observed in cursor rotation experiments.

### Experiments 1 to 3 – reaction times (RTs)

Reaction time (RT) has been shown to sharply increase at the onset of learning in visuomotor rotation tasks (Hinder et al. 2008; Shabbott et al. 2010; McDougle et al. 2015), and if RT is constrained, performance is significantly impaired (Fernandez-Ruiz et al. 2011; Haith et al. 2015) suggestive of strategy use. Figure 6 shows RTs from all experiments, summarized over 36 (18 for experiment 3) consecutive field trials (or channel trials for clamp) for selected blocks. In all experiments, RTs increased with the introduction of the force field: Across-block ANOVAs indicated significant differences for experiment 1 (*F*_*2.5,47.1*_=13.9, *p*<.001, *η*^*2*^_*gen*_=.42), and experiment 2 (*F*_*2.2,35.4*_=5.1, *p*=.009, *η*^*2*^_*gen*_=.24) and post-hoc t-tests confirmed significant differences from baseline to early field A (experiment 1: *t*=-4.1, *p*=.004; experiment 2: *t*=-3.1, *p*=.03). For experiment 3, ANOVA indicated a significant effect of block (*F*_*3.3,110.5*_=18.6, *p*<.001, *η*^2^_*gen*_=.35), but not of group (*F*_*1,34*_=.04, *p*=.85, *η*^2^_*gen*_=.001), and post-hoc t-tests showed a significant increase from baseline to early field A (*t=*-4.6, *p*<.001). By the end of field A practice, RTs were decreased relative to early field A in experiment 1 (*t*=5.5, *p*<.001), and 3 (*t*=7.2, *p*<.001), though this difference was not significant in experiment 2 (*t*=1.1, *p*=.6). RTs no longer differed significantly from baseline at the end of field A training (experiment 1: *t*=-1.4, *p*=.36; experiment 2: *t*=-1.4, *p*=.6; experiment 3: *t*=1.3, *p*=1.0). The introduction of field B caused another increase in RT in experiment 1 (compared to late field A: *t*=-3.3, *p*=.02), but not experiment 3 (*t*=.2, *p*=1.0). The latter fact may indicate that the highly variable behavior in experiment 3’s B-phase was truly reflective of learners not adopting a new strategy in response to field B, though this remains speculative.

**Figure 6:**
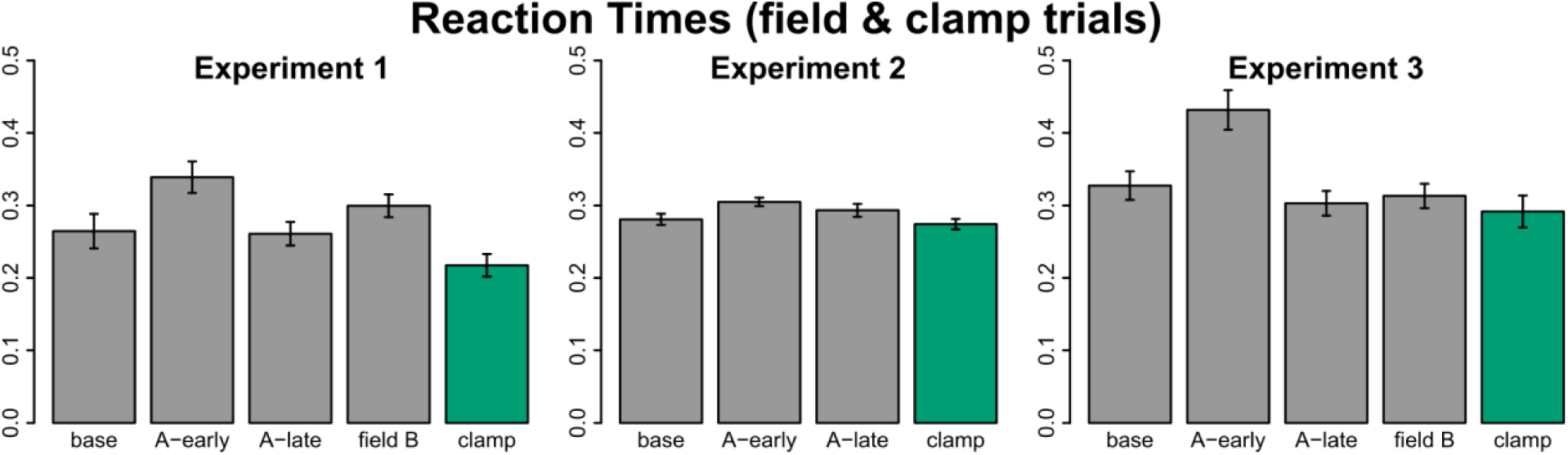
Across participant mean reaction times (±SEM) from selected blocks of 36 (experiment 1 and 2) or 18 (experiment 3) trials representing late baseline, early and late field A practice, complete field B practice, and early clamp, respectively. Grey bars are field trials, green bars are No Push trials for experiment 1 and 2 and right hand channel trials for experiment 3.

Critically, a purely implicit learning mechanism would not predict any measurable modulation of RTs upon the imposition of a perturbation. RT, however, is a reliable reflection of action selection/decision-making processes; thus, the observed RT increases likely reflect planning behaviors. After many trials of training, RTs are reduced, perhaps reflecting the “caching” of reinforced actions (McDougle and Taylor 2019), or habituation (Huberdeau et al. 2017; Hardwick et al. 2018). Importantly, whereas our interventions only tested for an explicit component on channel trials, the RT results suggest that this component also contributed on the standard force field learning trials.

## Discussion

We set out to test if an explicit learning component contributes to force field adaptation, similar to the way aiming strategies contribute to adaptation to cursor rotations (Heuer and Hegele 2008; Taylor et al. 2014). Our data support this notion in three ways. First, in experiment 1 and 2, instructing participants to not expect the force field on an upcoming (channel) trial reduced force expression compared to unannounced Catch channels. Conversely, instructing them to expect the force field caused no such reduction, showing that this was not just a stereotyped response to changes in trial conditions, but truly reflects the semantic content of the instruction. Despite the problems surrounding verbalization as a necessary criterion, responsiveness to the content of verbal instruction is a hallmark of explicit learning, whereas implicit adaptation is believed to be immune to this content (Mazzoni and Krakauer 2006; Heuer and Hegele 2008; Taylor and Ivry 2011; Schween et al. 2014). Second, in experiment 3, participants were able to express a learned pattern of compensatory forces with their left hand in response to a field learned with their right hand. This shows that they were both explicitly aware of the force field experienced with their right hand, and were able to intentionally utilize that knowledge to shape the motor response of their left hand. In addition to its intentional expression, our inference that left hand reports were qualitatively different from implicit adaptation is supported by the fact that participants appeared to report in different reference frames, which can be explained by differing interpretations of the instruction to “mimic the force”. Furthermore, right hand channel force displayed velocity dependence that increased with practice, whereas left hand force depended equally on position and velocity components. Velocity-dependence is a hallmark of adapting putative internal models to viscous force fields, and its reduced presence in the left-hand reports supports our interpretation that these reflect a qualitatively different component of learning. Third, across all experiments, introduction of the force field led to a transient increase in RT, in line with predictive compensation on normal field trials initially requiring some degree of cognitive processing, and subsequently becoming more proceduralized over time (Huberdeau et al. 2017; Hardwick et al. 2018; Leow et al. 2019; McDougle and Taylor 2019). Conversely, the introduction of the final clamp phase led to a reduction in RT, consistent with strategies being abandoned.

We take our results to show that some fraction of force field adaptation likely qualifies as volitional compensation. We find signs of this component contributing to predictive compensation on Catch channels, as well as to compensation on field trials. This raises one important question: Under which specific circumstances does explicit compensation operate, and how would this affect conventional interpretations of results from force field studies? We note that our experimental conditions were specifically designed to facilitate strategy formation and expression. Specifically, we instructed participants about the potential use of strategies. We considered this necessary as we suspected that reducing clarity in our instructions to test more “natural” behavior would have come at the risk of masking an explicit component through misinterpretation of instructions, particularly given the limited quality of explicit knowledge outlined above. For the same reasons, all of our experiments used only a single target direction, as this reduces the participants’ experience of the force field to mainly the horizontal component. Furthermore, we used rather large field constants in experiment 3; lower field constants would likely spur less explicit learning, similar to the way smaller or gradually introduced cursor rotations do (Malfait and Ostry 2004; Kagerer et al. 2006; Werner et al. 2014; Gaffin-Cahn et al. 2019). It therefore seems likely that the explicit component plays less of a role in force field paradigms that use small field constants, multiple targets, and deliberately instruct their participants to “not think about the task” (Morehead et al. 2017). In addition, velocity criteria that participants are frequently required to match may further draw attention away from strategic compensation and incentivize incidental learning (Lee et al. 2018b), as meeting them is relatively unrelated to reaching the target in a closed-loop movement.

Nevertheless, force field studies tend to vary in the magnitude of force applied, in the number of targets, and reaching characteristics, which, in turn, may vary the degree to which strategies are useful in compensating for the force. This may explain a number of mixed findings, such as savings, consolidation, and interference from declarative memory. The fact that we found signs of explicit learning across open- and closed-loop experiments and small and large field constants, respectively, indicates that the contribution of this volitional component is worth considering across experimental designs, particularly in situations where standard implicit adaptation may be insufficient.

A key issue here are situations where participants have to switch between different force environments. For example, when exposure alternates between two opposing force fields there is interference, leading to no learning on average (Gandolfo et al. 1996). Studies have identified contextual cues that allow learners to overcome this interference (Osu et al. 2004; Hirashima and Nozaki 2012; Howard et al. 2012, 2013, 2015; Nozaki et al. 2016; Sheahan et al. 2016; Heald et al. 2018; Proud et al. 2019). The relevant implication of our results here is that learners may in principle be able to learn such opposing force fields explicitly. Therefore, dual adaptation in many of these scenarios may reflect the formation of separate instances in declarative memory that support alternating compensation strategies (Gershman et al. 2014), rather than, or possibly prior to, the formation and adaptation of separate internal models (Wolpert and Kawato 1998). This might, for example, explain conflicting findings on the role of color cues in supporting dual adaptation (Gandolfo et al. 1996; Osu et al. 2004; Howard et al. 2013): their effect may be restricted to a strategic component whose contribution to solving the task at hand depends on certain experimental conditions. On a related note, switching between explicit strategies for a perturbed and a null condition using errors as a contextual cue has been suggested to explain savings in cursor rotation paradigms (Morehead et al. 2015). Our results imply that this explanation may also pertain to savings in force field adaptation.

As such, we propose that studies involving adaptation to force fields should be designed with the possibility of alternative learning mechanisms in mind, such as the explicit component described here. Notably, one may be tempted to consider explicit contributions an undesirable confound that should be eliminated; but it may also be true that they reflect an important stage in the learning of novel skills (Fitts and Posner 1967; Willingham 1998; Taylor and Ivry 2012; Stanley and Krakauer 2013). Thus, while it may be appropriate to limit explicit components in experimental manipulations where they would confound inferences about implicit learning, it may also be appropriate to embrace their existence, measure them, and further investigate their contribution to motor learning and control.

The notion that the explicit component may become particularly relevant upon switching between different perturbations also extends to the widely acknowledged idea that multiple timescales contribute to adaptation, which was originally inferred from behavioral phenomena that occur upon such switching (Smith et al. 2006). We have argued previously that “the fast timescale” of force field adaptation may be equivalent to an explicit learning component (McDougle et al. 2015). Whereas B-phase data in experiment 3 were too variable to provide further insight, experiment 1 is generally in line with this interpretation: Push and Catch channels, which include the explicit component, showed rapid adaptation to field B upon its introduction, whereas No Push channels, which exclude it, adapted considerably more slowly. Nevertheless, we note that not all aspects of the behavior we observed can be explained by a two-state model with one fast and one slow component. Crucial in this respect is that rebound occurs gradually. Since we instruct our participants to abandon the strategy at the beginning of the clamp phase, the two-state model would expect the explicit state to become disengaged and for behavior to abruptly switch to reflect only the implicit state, which should be in field A-direction and decay from there. Thus, whereas No Push channels during initial clamp not being at the same level as No Push channels in late field B can be explained by generalization of implicit learning being locally tied to the explicit movement plan (Hirashima and Nozaki 2012; Day et al. 2016; Sheahan et al. 2016; McDougle et al. 2017; Schween et al. 2018), the gradual drift that we observe at the beginning of the clamp phase indicates additional properties of underlying learning, such as a combination of two implicit and one explicit timescale (Miyamoto et al. 2014), or a gradual attenuation of strategic aiming rather than an immediate one.

In this manuscript, we have tended to contrast adaptation of internal models with explicit strategies. While we indeed believe that explicit strategies are qualitatively distinct from canonical adaptation of internal models and rely on different neural substrates (Anguera et al. 2010, 2012; Werner et al. 2014; Thürer et al. 2016), we would like to emphasize that this does not mean that explicit strategies do not rely on internal models or the cerebellum (Butcher et al. 2017). Internal models may serve various purposes, such as simulating potential outcomes to support (explicit) action selection (Barsalou 1999) and further research is needed to understand these relations.

In conclusion, our results show that a component of force field learning shares important properties with explicit aiming strategies in visual perturbation paradigms. It seems likely that the amount they contribute depends on specific experimental conditions, and more research is needed to clarify under which conditions these strategies are used and if they provide relevant alternatives to current explanations for behavioral observations. As a step to facilitate this clarification, we suggest that researchers should take care to report detailed information on instructions and other experimental conditions that may influence the generation and use of strategies.

## Acknowledgements

We thank Chandra Greenberg and Krista Bond for their support with data collection, as well as Peter Butcher, Carlo Campagnoli and Eugene Poh for their support with experiment setup. We also would like to thank Aaron Wong for help with the design of experiment 3 and Marion Forano for providing feedback on a previous manuscript version.

## Author contributions

RS, SDM and JAT conceived and designed research; RS, SDM, and JAT collected data; RS, SDM and JAT analyzed data; RS, SDM, MH and JAT interpreted results of experiments; RS prepared figures; RS drafted manuscript; RS, SDM, MH and JAT edited and revised manuscript; RS, SDM, MH and JAT approved final version of manuscript

---

Supplementary text 1: Scripted instructions of experiment 2 (experiment 1 was similar). As a general rule, we use this script as a guideline, but administer instructions in free speech with individual variations and answer questions asked by the participants within certain limits. While just reading the scripted instructions out loud or having participants read them may seem like better standardization, we consider our method more suited to achieve similar understanding on the side of participants rather than just giving them the same input.

## General task instructions (Instruct and demonstrate)

Welcome to our experiment. Today, we investigate how you learn to reach towards a target.

We will ask you to sit down in front of this robotic arm, grasp its handle and look into the screen below your eyelevel.

On each trial, the robot will guide your hand to a start location. Once your hand is in the start location, you will see a Plus-sign which I ask you to fixate with your view. After a short interval, the sign will disappear and a white target dot will appear. You don’t need to fixate on the Plus after it has disappeared.

Your task is to “shoot” through the target with the cursor in a swift, accurate movement. If you hit the target, it will turn green and you will hear a tone. If you miss it, neither of the two will happen. Independent of hit or miss, you may see a “Too Slow!” warning after the movement indicating that you should move faster. Your goal is to achieve the hit (i.e. target turning green and tone) without too slow warning as often as possible.

Note that the interval of target appearance is randomized – we do that so that you really have to react to the visual stimuli and cannot just fall into a rhythm. If you move before the target has appeared, the robot will resist your movement and bring you back towards the start location.

We will find you a comfortable position now. Please keep this position and take particular care not to move the chair.

## Instruct at Trial 90 (close to end of baseline)

After a few trials, the robot is going to push you off your path while you are moving towards the target. Please try to successfully hit the target in time as often as possible despite being pushed off!

Before some trials you will see a message instead of the Plus-Sign. If the message says “Push!” please expect the robot to push you off and act as on those trials. Note that you will not feel the robot push you on the trials with the “Push!” message but we nevertheless ask you to act as if it was going to! If the message says “No Push!” we would like you to act as in trials where the robot doesn’t push you off path and just move towards the target.

To sum up, please try to hit the target in time and stay alert to the messages. The important things for you to remember are that you treat “Push!” message trials as if the robot was going to push you even though you will not feel it, and that on “No push!” trials, act as if it was not going to push you.

Note that it is a hard task, so don’t be frustrated if you don’t immediately manage to hit. Try your best and see if you can get better.

## After trail 550 (before Error Clamp)

Starting now, for the remainder of the experiment, we would like you to not push deliberately or apply any aiming strategies you may have developed. Just move towards the target. This is the same as on “No push!” trials, but you won’t be seeing any more messages.

